# Hemispheric Dissociation Revealed by Attentional Isolation and tRNS

**DOI:** 10.1101/2025.06.17.660112

**Authors:** Michele Tosi, Giulia Ellena, Federica Contò, Grace Edwards, Lorella Battelli

## Abstract

Prolonged sensory imbalance, induced by directing attention to one visual field, can paradoxically enhance performance in the opposite, non-attended visual field. This effect is likely driven by the brain’s homeostatic mechanisms that regulate excitation and inhibition between hemispheres in homotopic attention processing regions. Here, we employed transcranial random noise stimulation (tRNS) to modulate cortical excitability and probe its role in interhemispheric dynamics controlling visual attention. Specifically, we used a procedure called attentional isolation, where neurotypical participants covertly focused their visual attention in one hemifield (the attended visual field) for 30 minutes. Performance changes in both the unattended (opposite) visual field and the attended visual field were measured following this manipulation. We applied transcranial random noise stimulation (tRNS) over the right or left frontoparietal cortex to modulate the excitability of one hemisphere relative to the other during attention isolation, probing the neural mechanisms underlying the observed contralateral performance shift. Our results showed improved performance in the previously unattended visual field following the attentional isolation period after sham stimulation. However, tRNS revealed a functional dissociation between the hemispheres: right hemisphere active stimulation abolished the performance improvement, while left hemisphere stimulation preserved it. These findings suggest distinct roles for the left and right hemispheres in modulating paradoxical visual performance shifts and may inform the development of novel neurorehabilitation strategies for clinical populations.

## INTRODUCTION

Our brain has limited computational resources and must constantly select behaviorally relevant information to prevent saturation (Nobre & van Ede, 2023). It must thus prioritize relevant information while simultaneously suppressing irrelevant ones, and selective attention plays a central role in this process (Carrasco, 2011, Kastner & Ungerleider, 2000, Desimone & Duncan, 1995). Neurophysiological and imaging evidence indicate that visuospatial attention relies on broad functional networks involved in top-down and bottom-up attentional mechanisms. In particular, the dorsal attention network (DAN) is the key cortical network controlling endogenous focus on goal-relevant attentional mechanisms (Corbetta & Shulman, 2002). Further studies have shown that the strength of the functional connectivity between different nodes of the DAN is associated with behavioral improvement in visuospatial tasks (Contò et al., 2021; Machner et al., 2022).

Normal behavioral attentional functioning is maintained by an intracortical balance mediated by excitation and inhibition between homologous areas in the left and right hemispheres via transcallosal connections (Bertolucci et al., 2018; Cazzoli et al., 2009; Corbetta et al., 2005; Giglia et al., 2011; Sparing et al., 2009; Tatti et al., 2017). Neurophysiological studies have shown that these connections almost entirely consist of excitatory fibers, and only during simultaneous bilateral stimulation can inhibitory mechanisms arise (Palmer et al., 2012). Alterations of inter-hemispheric balance are commonly associated with a wide range of neurological and psychiatric conditions, such as Huntington’s disease, autism and schizophrenia (Ben-Ari et al., 2012; Cummings et al., 2009; Ridding et al., 1995; Rubenstein & Merzenich, 2003; Vierling-Claassen et al., 2008). A key concept of the mechanisms maintaining intracortical balance between the two hemispheres is mutual inhibition (Beaulé et al., 2012, Carson, 2020, Zebhauser et al., 2019). Current literature emphasizes the role of inter-hemispheric inhibition not only as a mechanism to prevent over-excitation but also as a means to modulate the neural output of specific circuits that are segregated and lateralized in the brain (Carson, 2020). As a consequence, a stroke-induced disruption of this mutual inhibition between the two hemispheres, especially in the parietal areas, is commonly associated with hemispatial neglect, a deficit that leads to impairments in visuospatial performance within the contralesional (neglected) hemifield (Casula et al., 2021). Interestingly, the degree of disturbance in the excitation/inhibition balance can correlate with the severity of the neglect symptomatology (Baldassarre et al., 2014; Koch et al., 2013). One way to partially restore and modulate the balance between excitation and inhibition is through the application of noninvasive brain stimulation, which has demonstrated potential for recovering visual attentional abilities in neglect patients (Agosta et al., 2014; Smit et al., 2015; Nyffeler et al., 2019).

The modulatory effects of noninvasive brain stimulation can also help understand the interhemispheric interactions in a healthy brain. Specifically, transcranial random noise stimulation (tRNS) has been successfully used to modulate cortical excitability (Herpich et al., 2018) and affect functional connectivity within and between brain networks in healthy volunteers (Contò et al., 2021). It has also yielded promising results in the motor (Moret et al., 2019; Qi et al., 2019, Ellena et al., 2025), visuospatial (Contò et al., 2023), visuoperceptual (Camilleri et al., 2016; Herpich et al., 2019), mnestic (Murphy et al., 2020) and auditory domains (Van Doren et al., 2014). tRNS consists of the application of a weak alternating current at random frequencies and fixed intensity to the scalp. Although its physiological mechanism is not fully understood, one hypothesis suggests that tRNS enhances neural activity by increasing the signal-to-noise ratio, which may contribute to improved performance in perceptual and cognitive tasks (Fertonani et al., 2011; Antal & Herrmann, 2016; van der Groen et al., 2022).

The concept of intracortical balance has also been explored in studies investigating the relationship between neural imbalance and sensorimotor improvement (Andrushko et al., 2023) as well as in research on visual detection (Binda et al., 2018; Lunghi et al., 2015). These studies highlight the role of GABA (gamma-aminobutyric acid) as an inhibitory neurotransmitter involved in regulating excitation/inhibition balance in experience-dependent cortical plasticity. For example, Andrushko et al. (2023) demonstrated that unilateral handgrip training led to improved motor function in the non-trained hand, a phenomenon known as cross-training (or cross-education) with significant implications for neurorehabilitation (Ehrensberger et al., 2016; Green & Gabriel, 2018; Kay et al., 2024; Lim & Madhavan, 2023). A further example of the brain homeostatic regulatory mechanisms is described by Lunghi et al (2015). Specifically, they described that a temporary visual deprivation manipulation led to an increase in neural responsiveness in the deprived eye, with GABA-mediated changes playing a crucial role. These findings suggests that prolonged sensory imbalance can trigger compensatory neurophysiological adaptations aimed at restoring homeostatic balance (Boroojerdi et al., 2001; Frangou et al., 2019; Mrsic-Flogel et al., 2007;Wenner, 2011).

In the present study, we aimed to investigate the neurophysiological mechanisms underlying the interhemispheric interactions in attentional control. Using *attentional isolation*, a paradigm developed by Edwards et al. (2021), we induced a temporary imbalance in visuospatial attention by requiring participants to attend to one hemifield for an extended period. Edwards et al., showed enhanced performance in the previously unattended visual field following a period of attention isolation, suggesting this behavioral manipulation modulates excitatory-inhibitory interaction between homotopic attention processing regions. Here, we will concretely explore the excitatory-inhibitory mediation of *attention isolation* with the addition of tRNS over frontoparietal areas corresponding to the frontal eye field (FEF) and the intraparietal sulcus (IPS) regions implicated in attentional selection (Xu & Jeong, 2015) and endogenous attentional shifts (Heinen et al., 2017 Thompson et al., 2005). By modulating cortical excitability, we aimed to assess how neural plasticity mechanisms contribute to the observed behavioral effects.

As right-hemisphere attention processing regions play a dominant role in visuospatial attention (Bartolomeo & Seidel Malkinson, 2019; Brancaccio et al., 2022), we expected that tRNS over the right frontoparietal cortex would interfere with the beneficial effects of attention isolation. Specifically, given the right hemisphere’s field-wide representation and its integrative function across both hemifields, increasing its excitability may disrupt the interhemispheric imbalance necessary for attentional reallocation. In other words, we hypothesized that enhancing right frontoparietal excitability during unilateral attention would reduce the neural contrast between attended and ignored hemifields, thereby suppressing the compensatory behavioral gain typically observed in the unattended field. This prediction stems from the idea that right-lateralized stimulation may unify attentional processing across the visual field, counteracting the homeostatic mechanisms triggered by isolation.

Conversely, left hemisphere attention processing regions are considered to be more localized to contralateral attentional processing (Sheremata & Silver, 2015; Duecker et al., 2013). Therefore with tRNS to left frontoparietal regions we expected preserved or even enhanced behavioral effects of attentional isolation. By increasing excitability in a network that primarily modulates attention to the right visual field, left-sided stimulation should allow the right hemisphere to continue regulating the contralateral hemifield, maintaining the interhemispheric asymmetry necessary for compensatory gain. In this way, left hemisphere stimulation may support the segregation of visual fields— facilitating rather than disrupting the plasticity induced by unilateral attention. These predictions allowed us to test whether hemispheric stimulation differentially integrates or isolates attentional control across the visual field.

By delineating these hemispheric dissociations, we aimed to provide new insights into the neural mechanisms underlying attentional plasticity and to inform future neurorehabilitation strategies for clinical populations.

## Experiment 1 – Unilateral *right* stimulation

### Methods

#### Participants

Forty-five neurologically healthy (mean age 24.83, SD 3.84, 26 female), right-handed individuals, with normal or corrected-to-normal vision were enrolled in this study. All subjects met all brain stimulation screening criteria (Fried et al., 2021) and provided informed consent approved by the ethics committee of the University of Trento. The study conformed to the Declaration of Helsinki. Subjects received monetary compensation for their participation in the study. Six subjects after the first session were excluded from the experiment due to poor compliance with the study design, leaving a total of 39 subjects. Poor compliance was primarily attributed to difficulties in adjusting to the eye-tracking acquisition and maintaining central fixation for a prolonged period of time.

#### Bilateral multiple object tracking

Participants performed a bilateral multiple object tracking (MOT) task, which requires participants to track two moving disks in each hemifield among distractors. In each trial, four disks were displayed in each hemifield within a 6 cm by 6 cm area (four disks on the right hemifield and four disks in the left hemifield). All disks were black (Figure 1B), with a radius of 0.25cm. Each disk could never cross the midline and repelled one another to avoid collisions. Two disks on each side of the fixation briefly flashed at 4 Hz, indicating the target disks to track during each trial. Importantly, the selection of target disks was random, and participants could not predict in which visual field they would be tested. Following the flashing cues, all the disks moved at a constant speed for duration of 2 sec. The individual speed threshold for each participant was measured at baseline using a staircase procedure (see the “Thresholding MOT” section for details). At the end of the 2-second movement period, all disks stopped, one disk turned red and subjects were asked to indicate whether it was a target or a distractor by entering their response on a keyboard. The target could appear randomly on the right or in the left visual field. Response feedback was provided, with the fixation point turning green for correct and red for incorrect responses. Throughout the task, participants were instructed to maintain their gaze on the fixation, monitored with an eye tracker (see “Eye-tracking acquisition” section for details). When they broke the fixation, the trial was aborted, and a new trial began. On average, participants performed 106 trials on pre-test (SD = 6.61), 309 on attentional isolation (SD=25.45) and 107 on *post*-test (SD=6.23). During the 30 minutes of attentional isolation, on average subjects performed 103 trials (SD=9.58) every 10 minutes. Participants were tested on a Samsung 2233RZ LCD (22”) monitor (screen resolution 1680×1050), at a distance of 70 cm. The experiment was run though a 2020 HP Pavilion Gaming Laptop (Windows 10 installed). The stimuli were presented using the psychophysical toolbox PsychoPy 2022.2.0.

**Figure 1.**
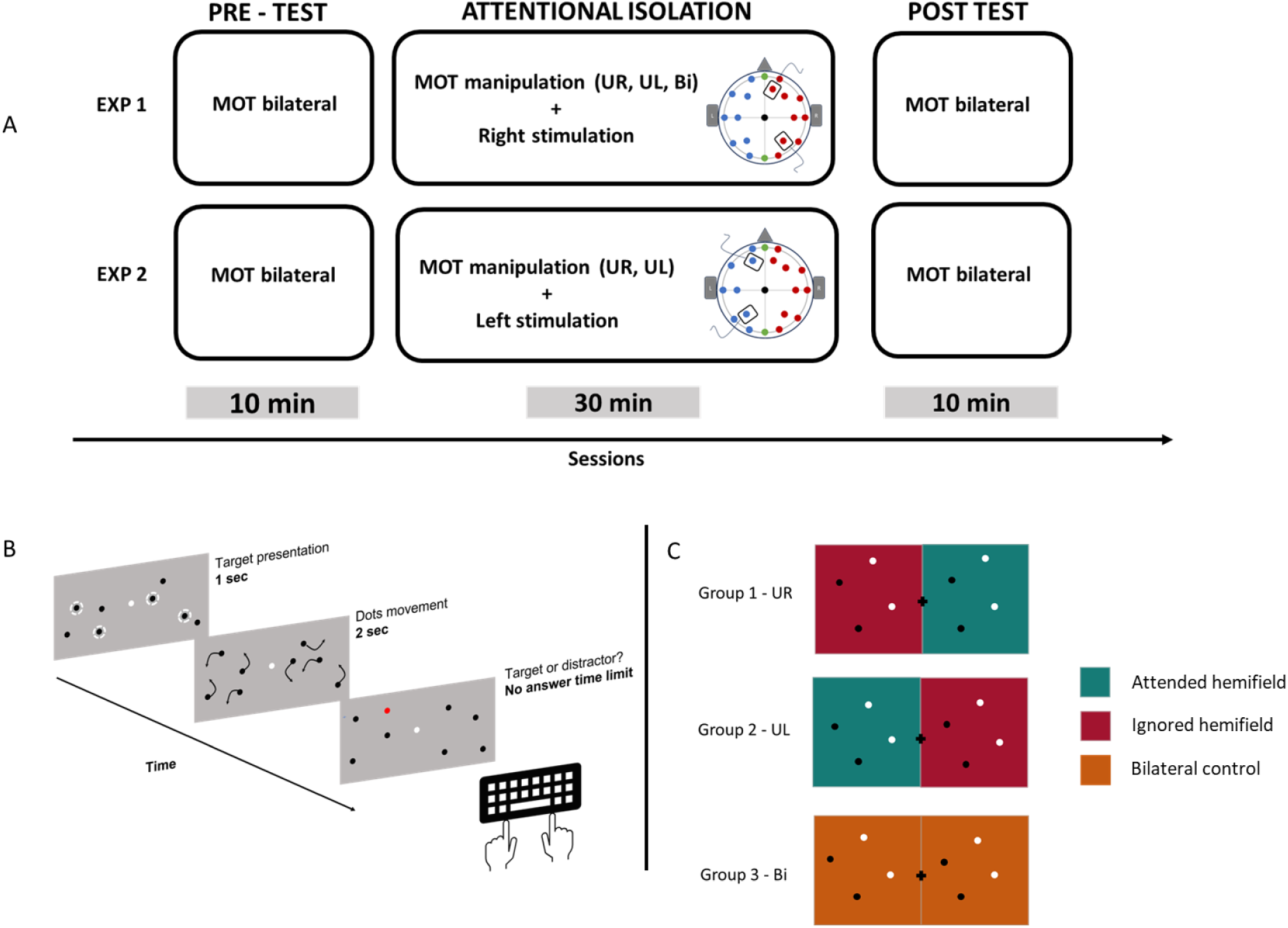
Procedure and stimulation locations. **(A)** At the beginning of the experiment, threshold and pre-test performance were assessed. Subjects were then assigned to a specific “isolation group” (UL - unilateral left, UR - unilateral right, Bi - bilateral control). The attentional isolation lasted 30 minutes. Simultaneously, from the beginning of the attentional isolation, subjects received 20 minutes of tRNS at 2mA. After the manipulation, subjects immediately underwent bilateral post-test and a retest 10 minutes after the end of the post-test. **(B)** MOT design - This task involved identifying and tracking of specific objects (two moving disks) among a group of distractor disks. **(C)** Hemifield tracked during the 30 minutes of attentional isolation for each group. Each color represents one tracking condition.

#### Study design

Experimental design is briefly summarized in Figure 1A. Subjects were randomly assigned to three different manipulation groups: Group 1 performed a right unilateral MOT, Group 2 performed a left unilateral MOT, and a control Group 3 underwent a bilateral MOT task (Figure 1C). The manipulation time lasted 30 minutes for each session. During unilateral MOT, participants were instructed to track two moving target-disks among two moving distractor-disks presented in one hemifield only (a total of 4 disks). Four additional moving disks were simultaneously displayed in the contralateral hemifield to maintain a homogeneous sensory input between the two hemifields. During the bilateral MOT control condition, participants were asked to simultaneously track four target disks (two in each hemifield) among four distractor disks (two in each hemifield). Participants were required to focus on the attended hemifield while actively ignoring the unattended contralateral hemifield. The task was gaze-contingent (see section *Eye-tracking acquisition)* and when subjects broke fixation the trial was aborted, and a new trial started.

All Subjects (Group 1, 2 and 3) underwent two counterbalanced sessions: a sham and an active stimulation condition separated by at least seven days. In Experiment 1, 39 subjects received stimulation over the right hemisphere. While, in Experiment 2, 26 subjects received stimulation over the left hemisphere. Based on previous literature (Edwards et al., 2021) and on the results of Experiment 1, we decided to not include a bilateral MOT condition in Experiment 2, since we already expected no effect in this control condition. For both experiments, 13 subjects were assigned to each group.

#### Thresholding MOT

At the beginning of each experimental session, we psychophysically measured participants’ speed threshold in the MOT task. A two-alternative forced choice (2AFC) with fixed step sizes staircase was employed to determine the speed at which subjects performed at 70% correct. We used a staircase procedure to adjust the stimuli motion speed based on participants’ performance: it increased by 0.5 cm/sec after two correct consecutive responses and decreased after one single incorrect response. The staircase procedure continued until it reached a predefined endpoint of 20 reversals. Importantly, only the data from the last six reversals were used to estimate individual accuracy performance, ensuring a stable and reliable measure of participants’ tracking abilities before the experimental manipulation. Prior to the thresholding, subjects practiced bilateral MOT for 120 sec, at a fixed speed of 3.5 cm/s. On average, participants performed the MOT after the threshold with a mean speed of 8.17 cm/sec, std 2.56 cm/sec, ranging from 2.5 cm/sec to 15.17 cm/sec.

#### Eye-tracking acquisition

Central fixation was continuously monitored with an EyeLink 1000 Plus eye-tracking system (SR Research). Calibration and validation were performed at the beginning of each run. Throughout the experiment, a box of 3° × 3° was placed around the central fixation point. If participants shifted their gaze outside of this boundary box, the trial would restart.

#### Stimulation protocol

A battery-powered device (DC-Stimulator, manufactured by NeuroConn in Ilmenau, Germany) was used to deliver the electrical stimulation. Electrodes, each measuring 5×5 cm² and inserted in sponges soaked with saline solution, were positioned according to the 10/20 International electroencephalographic system (Herwig et al., 2003; Okamoto et al., 2004), over F2 and P4 for the right and F1 and P3 for left hemisphere, roughly corresponding to the FEF and the IPS, in each hemisphere. In Experiment 1, electrodes were placed over the right, while in Experiment 2 over the left hemisphere. The stimulation was administered for the first 20 minutes within the period of isolation, with a current intensity of 2mA peak-to-peak at a high frequency range ranging randomly from 101 to 640 Hz. A gradual fade-in/fade-out period lasting 10 sec preceded and followed the stimulation. In the sham condition, the stimulation ceased after 30 sec plus the fade-in/fade-out period. Importantly, participants did not report any significant adverse effects following the stimulation.

#### Data analysis

Statistical analyses were conducted in RStudio, using the *tidyverse* and *lme4* packages (Bates et al., 2014; Wickham et al., 2019). Figures were made in R package *ggplot2* (Wickham H, 2016). We first checked if the pre-test baseline performance was equal across groups. Pre-test performances were tested with a specific *Generalized Linear Mixed-Effects Models (glmer)* from the *lme4* package. The between subject’s predictors were Manipulation (attended visual field, ignored visual field, or control) and Montage (right montage, left montage); the within subjects’ predictor was visual field (left, right), stimulation (sham, active) and Session (Pre,Post). Outliers within visual field per each experiment at baseline were removed with the interquartile range methods. Specifically, average individual performance values that surpassed the upper quartile plus 1.5 times the interquartile range (IQR) or fall below the lower quartile minus 1.5 times the IQR, were identified as outliers and removed from the analysis. Analysis for detecting changes in performance from pre to during to post were ran with a *glmer* model. Post-hoc comparisons were performed through *emmeans* package (Lenth R, 2024), applying the *Holm* correction method. Finally, the comparison of gain in performance (delta, calculated as post - pre) was performed using a Linear Mixed-Effect model (*lmer*).

### Results

#### Isolation manipulation impact on tracking performance

We first checked for outliers in the pre-test conditions, and we did not find any for either the *left* or *right* visual field. Importantly, we did not find any *pre*-test baseline difference in performance between manipulation conditions (χ2(2) = 0.30, p = 0.85, *glmer*) and stimulation conditions (χ2(1) = 0.64, p = 0.42, *glmer*). First, we investigated the impact of isolation manipulation on tracking accuracy (attentional gain) in the ignored visual field relative to the (bilateral tracking) control condition. We expected an increase in tracking accuracy in the ignored visual field after the manipulation period and no significant changes from pre to post in the attended visual field and in the bilateral tracking condition. We then examined the effect of right frontoparietal stimulation on attentional gain following the manipulation period. We found a three-way interaction between manipulation, session, and stimulation, suggesting that the effect of the manipulation changed across sessions and between stimulation conditions (χ2(2) = 6.69, p = 0.03, *glmer*). A post-hoc comparison revealed a change from pre- to post-manipulation in the ignored visual field in the sham condition (estimate = 0.48, se= 0.08, z=5.45, p<.01, Figure 2 - sham), while we did not find any change in performance between the pre- and post-manipulation in the ignored visual field in the active stimulation condition (estimate = −0.05, se=0.08, z=-0.67, p>.05, Figure 2 - active). Therefore, attentional isolation resulted in a significant increase in tracking accuracy in the ignored visual field during sham. However, the significant gain in accuracy was abolished after right frontoparietal stimulation. Importantly, to see whether unilateral tRNS had any effect in the absence of any attentional manipulation, we also compared performance in the left hemifield, contralateral to the stimulation, in the bilateral control group for sham versus active stimulation. We found no change in response between active and sham (χ2(1) = 0.28, p = 0.59, *glmer*), indicating that tRNS had a significant impact only during the attentional manipulation.

**Figure 2.**
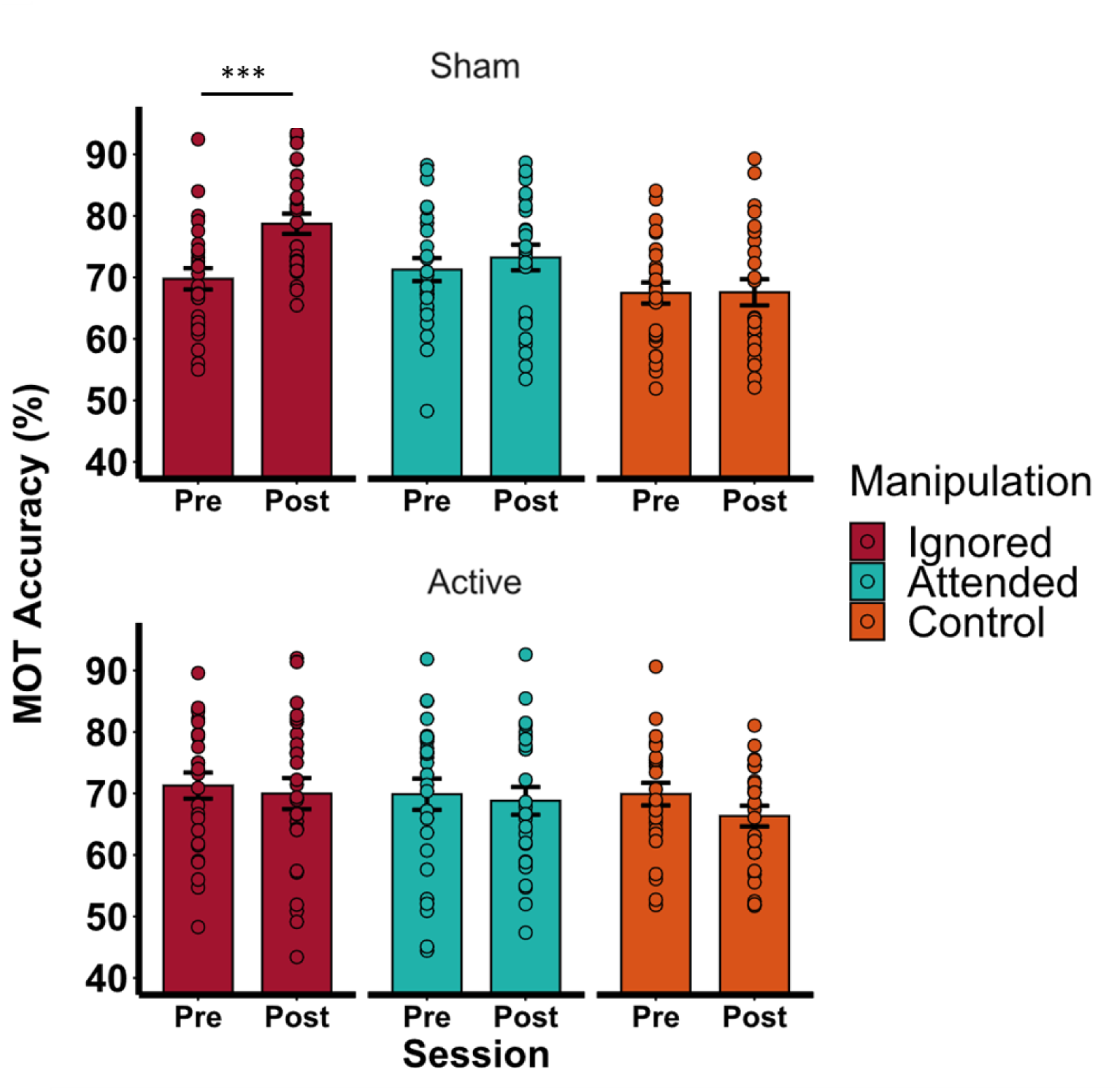
Change in accuracy from Pre to Post over different Manipulation conditions (Ignored hemifield, Attended hemifield, Control), for the Sham (top graph) and Active (bottom graph) stimulation for the right montage condition

## Experiment 2 – Unilateral *left* stimulation Methods

The procedure was kept identical to Experiment 1, except for the stimulation site. We also did not include a bilateral tracking control group as we already replicated the effect we found in our previous study and to increase the statistical power (Edwards et al., 2021). We used the same electrode montage as in Experiment 1, but for *left* FEF-IPS (corresponding to electrodes F1-P3) in accordance with the 10-20 international EEG system. Twenty-nine neurologically healthy (mean 22.27, std 2.65, 19 female) right-handed individuals, with normal or corrected-to-normal vision were enrolled in this study. Three subjects were excluded from the experiment after the first session due to poor compliance with the study design, leaving 26 subjects. All subjects met all brain stimulation screening criteria (Fried et al., 2021) and provided informed consent approved by the ethics committee of the University of Trento. The study conformed to the Declaration of Helsinki. Subjects received monetary compensation for their participation in the study.

### Results

#### Isolation manipulation impact on tracking performance

We first checked for outliers in the pre-test conditions for all subjects. After the outlier detection, four subjects were removed from the sample. Importantly, we did not report any pre-test differences neither between manipulation conditions (χ2(1) = 2.33, p = 0.12, *glmer*), nor between stimulation conditions (χ2(1) = 0.41, p = 0.51, *glmer*). Similar to experiment 1, we were interested in investigating the effect of a prolonged unilateral tracking period over the contralateral, ignored portion of the visual field. Furthermore, we expected left frontoparietal tRNS to have a weaker impact on performance compared to right frontoparietal stimulation. As previously reported, we expected an improvement in the ignored visual field compared to pre-test. We did not find a three-way interaction between session, manipulation and stimulation, (χ2(1) = 0.001, p>.05, *glmer*). However, we found a two-way interaction between session and manipulation, suggesting that the effect of attentional isolation changed across time, independently from the stimulation (χ2(1) = 11.18, p<.01, *glmer*). Post hoc analysis revealed a change in tracking accuracy from pre- to post-manipulation in the ignored visual field (estimate=0.26. se=0.06, z=4.04, p<.01; Figure 3), indicating a gain in performance over the ignored visual field. We also reported a change in performance from pre to post in the attended visual field (estimate= −0.17, se=0.06, z=-2.68, p<.05; Figure 3), revealing a detrimental effect on tracking accuracy over the attended visual field after the attentional isolation.

**Figure 3.**
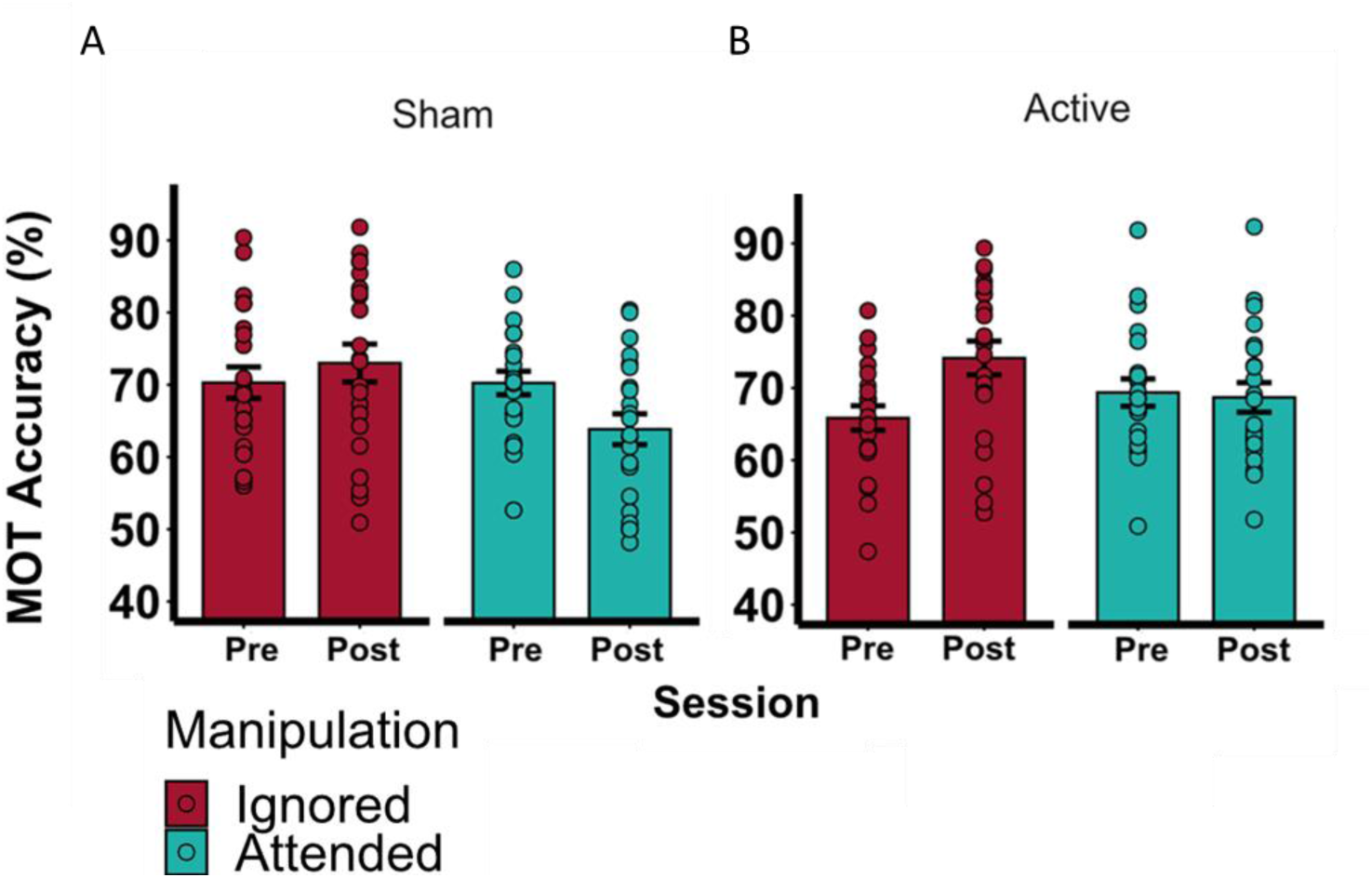
Change in accuracy from pre- to post-manipulation for the Sham (A) and for the Active (B) stimulation condition, for the Ignored (red) and the Attended (green) visual hemifield for the Left Montage stimulation condition. In our model, Sham and Active did not report any statistical differences.

#### Comparison between left and right hemisphere stimulation in the ignored visual field

Based on the results of Experiments 1 and 2, we decided to further analyze the changes in pre-versus post-stimulation performance between left and right stimulation in both experiments over the ignored visual field. We found a two-way interaction between montage and stimulation, indicating that the effects induced by stimulation varied between right and left montages (χ2(1) = 16.40, *p* < .01, lmer). As expected, we found a change in accuracy from pre- to post-manipulation (delta) in the ignored visual fields between the right and left (active) stimulation conditions (estimate= −0.09, se=0.03, t=-3.18, p<.05) indicating that the magnitude of attentional enhancement over the ignored visual field is different following left and right hemisphere stimulation, thus highlighting a dissociation in the stimulation effects between the left and right hemispheres (Figure 4). Finally, we found no differences in the magnitude of attentional enhancement in the ignored visual field in the sham condition in both Experiments 1 and 2.

**Figure 4.**
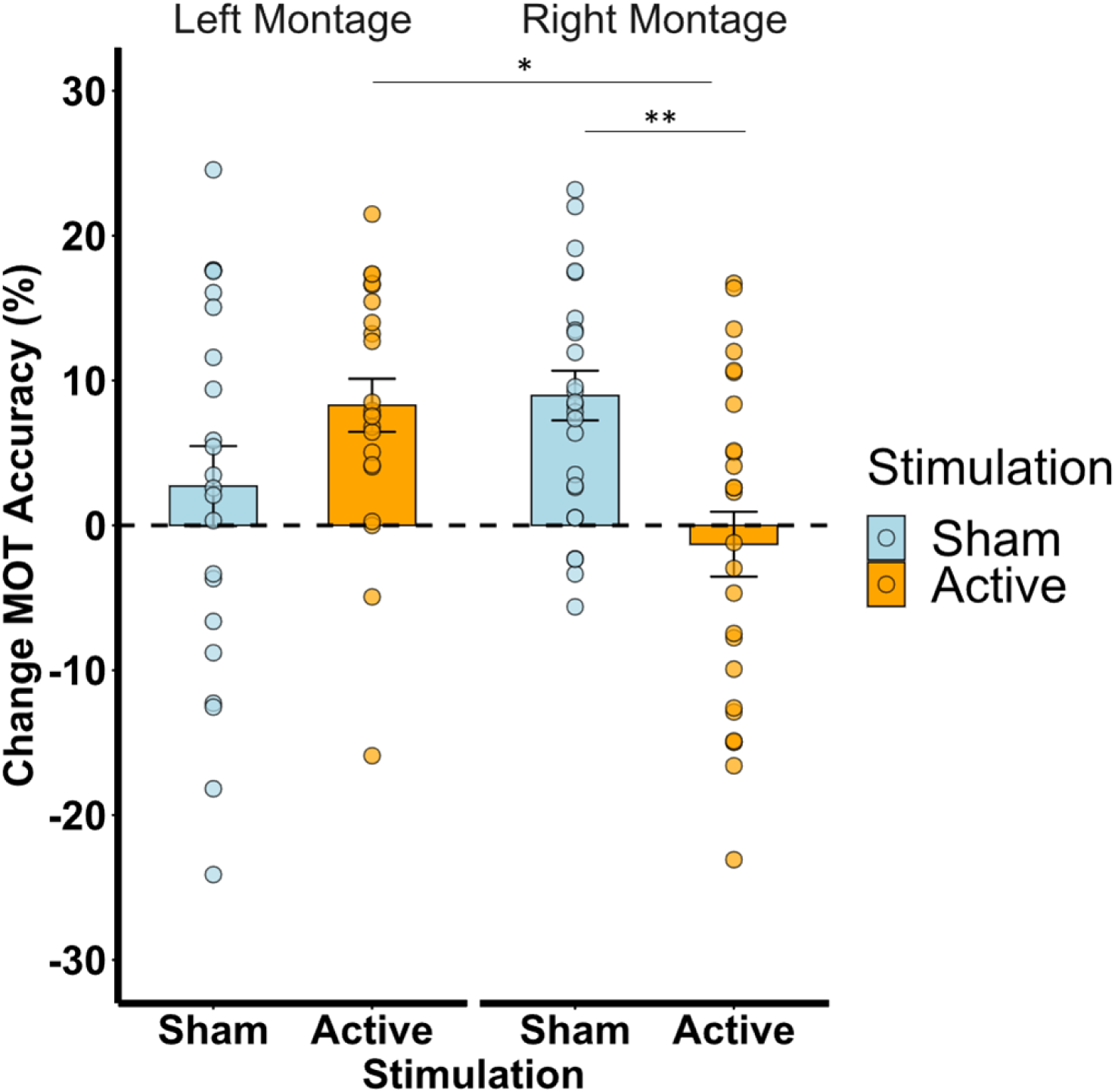
Changes in performance pre– versus post–stimulation for the right and left montage in the Ignored Visual Field.

#### Hemispheric-dependent response to tRNS during the attentional isolation

We conducted additional analyses to examine the effect of attentional isolation in the attended visual field to test for hemispheric asymmetries during the manipulation. Given that the beneficial post-manipulation effect was present after the left frontoparietal tRNS but was abolished after the right tRNS, we sought to better understand the dynamics of the cortical response to stimulation during the manipulation in the attended visual fields (isolation side). Specifically, we ran a model to examine changes in performance over the isolation side (ipsilateral or contralateral to the stimulation site) during the 30 minutes of attentional isolation in the active condition relative to sham for both hemispheres (left and right montage). We found a three-way interaction between stimulation, montage, and side of isolation (χ2(1) = 26.57, p <.01, *glmer*), suggesting that the effect of the stimulation (sham or active) during the attentional isolation manipulation depended on montage (left or right hemisphere) and the attended side (ipsilateral or contralateral to stimulation). Right stimulation during attentional isolation had a detrimental effect for both ipsilateral and contralateral sides (Figure 5, right montage). Specifically post-hoc analysis revealed that there was a decrease in tracking accuracy during right hemisphere stimulation relative to sham in the (attended) contralateral side (estimate = −0.23, se =0.05, z = −3.90 p<.01) and during right hemisphere stimulation relative to sham on the (attended) ipsilateral side (estimate =-0.36, se =0.05, z = −6.45, p<.01). This suggests that performance in both hemifields was impaired by the application of tRNS over the right hemisphere at the same time as the task. In other words, the effect of stimulation during the attentional isolation manipulation was detrimental bilaterally (Figure 5). Furthermore, we observed an improvement in ipsilateral tracking accuracy during left hemisphere stimulation compared to the sham condition (estimate =0.23, se =0.05, z = 3.91, p<.01), wheras contralateral tracking accuracy decreased during left hemisphere stimulation compared to sham (estimate =-0.25, se =0.06, z = −3.70, p<.01). Finally, we did not find an effect of the right stimulation on the bilateral control group during the attentional isolation (χ2(1) = 0.82, p > .05, *glmer*). In short, left hemisphere stimulation during task performance tends to facilitate tracking performance ipsilateral to stimulation, while contralateral tracking performance is impaired (Figure 5, left montage).

**Figure 5.**
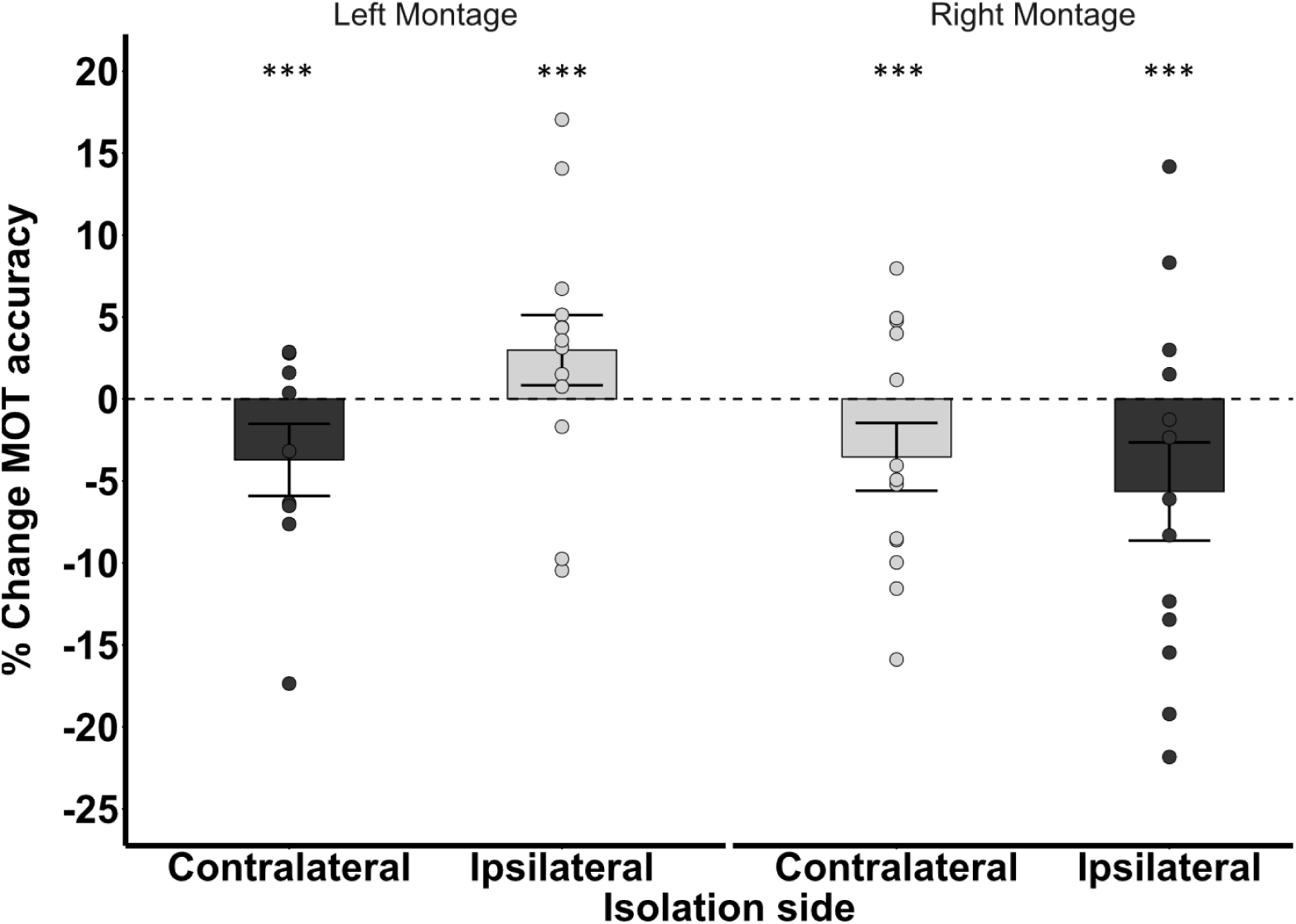
Percent change in MOT accuracy reported as difference between Active and Sham (delta) for isolation sides and condition.

### Discussion

Our results show a clear hemispheric asymmetry in how frontoparietal transcranial random noise stimulation (tRNS) modulates attentional reallocation. Specifically, left frontoparietal tRNS preserved the post-manipulation improvement in the ignored visual field observed *after* participants performed a unilateral multiple objects tracking task. In contrast, right frontoparietal tRNS abolished this beneficial effect observed in previous work (Edwards et al., 2021). *During* the manipulation period, when participants were asked to focus on tracking multiple moving objects in one visual hemifield for 30 min while actively ignoring objects in the opposite, unattended field, tRNS effects were also hemisphere-specific: right hemisphere stimulation impaired performance in both visual fields, whereas left hemisphere stimulation selectively reduced performance in the contralateral visual field and improved it ipsilaterally. These effects were not present in a bilateral multiple objects tracking control condition, suggesting that the effects were solely due to the combination of manipulation and stimulation.

These results suggest that left frontoparietal stimulation may facilitate adaptive attentional reallocation, while right hemisphere stimulation disrupts it, likely due to differences in each hemisphere’s role in spatial attention. The beneficial behavioral effect of attentional isolation observed in the sham condition replicated earlier findings (Edwards et al., 2021), but the concurrent application of active tRNS altered this outcome. In particular, right frontoparietal tRNS eliminated the post-manipulation gain, suggesting a disruptive interaction between stimulation and endogenous attentional processes during the manipulation. Thus, our results underscore the importance of hemispheric specialization in attentional control. The right hemisphere’s dominant role in spatial attention (Corbetta & Shulman, 2002) may explain why stimulation here leads to bilateral suppression of tracking performance (Hilgetag et al., 2001; Szczepanski & Kastner, 2013). In contrast, left hemisphere stimulation produced more localized, contralateral effects (Figure 5), consistent with prior reports (Battelli et al., 2009, 2017).

These findings align with the broader literature on hemispheric asymmetries: the right hemisphere exerts bilateral attentional control, while the left hemisphere shows a stronger contralateral bias (Heilman & Van Den Abell, 1980; Corbetta & Shulman, 2002). Furthermore, tRNS likely impacts the dorsal attention network, altering how attentional signals are integrated across hemispheres. Given tRNS’s effects on cortical excitability and connectivity (Chaieb et al., 2011; Contò et al., 2021), the observed behavioral changes suggest that these network-level modulations are sensitive to both the site of stimulation and the attended visual field.

Taken together, these findings indicate that unilateral tRNS may suppress attentional deployment across both fields during attentional isolation, particularly when targeting the right hemisphere, while left hemisphere stimulation can have facilitatory effects in the ipsilateral field (see Figure 5). This supports neuropsychological models of interhemispheric competition and attentional vectors (Bartolomeo, 2000, 2006; Chica et al., 2012; Kinsbourne, 1987, 1993; Mesulam, 1981; Vallar & Bolognini, 2014). In our study, right hemisphere tRNS during the manipulation may have induced inhibitory effects that disrupted attentional deployment bilaterally—consistent with its broad attentional influence (Shulman et al., 2010). Conversely, left tRNS impaired contralateral field tracking but improved performance in the ipsilateral field, likely due to a combination of lateralized attentional vectors and compensatory mechanisms from the dominant right hemisphere. This may explain why, in Experiment 2 (left stimulation), we observed preserved post-manipulation benefits, while in Experiment 1 (right stimulation), performance returned to baseline with no gain (Edwards et al., 2021).

From a clinical perspective, these hemispheric effects have potential implications. Frontoparietal stimulation’s impact on attentional asymmetries could inform neuromodulation strategies for patients with conditions such as hemispatial neglect (Shindo et al., 2006; Agosta et al., 2014). Tailoring stimulation protocols based on hemispheric roles may enhance rehabilitation outcomes, and these principles could extend to other attention-related disorders (Agosta et al., 2014; Edwards et al., 2019). The dominant role of the right parietal cortex in neglect is well-established (Bartolomeo & Chokron, 1999; Bisiach et al., 1986; Umiltá, 1995), and Kinsbourne’s interhemispheric competition theory provides a useful framework: both hemispheres direct contralateral attention and inhibit each other, with the left hemisphere biasing rightward attention. Damage to the right hemisphere—or its suppression via noninvasive brain stimulation—can result in reduced attentional deployment to the left field (Kinsbourne, 1987; Sparing et al., 2009).

Lateralized effects of tRNS observed here likely reflect distinct roles of the left and right frontoparietal networks. The right IPS and FEF are involved in global attention allocation and distractor suppression, critical in our task (Szczepanski & Kastner, 2013; Corbetta & Shulman, 2002). Disrupting this network with right tRNS likely reduced attentional capacity bilaterally during the manipulation, thus abolishing the post-manipulation benefit (Edwards et al., 2021). In contrast, left hemisphere stimulation selectively impaired right-field attention, perhaps due to compensation by the right hemisphere, while preserving the post-manipulation benefit (Hilgetag et al., 2001; Duecker et al., 2013).

Dynamic tracking requires stable excitability in lateralized networks (Howe et al., 2009). Disruptions in excitability, whether through stimulation or pathology, can inhibit contralateral processing, as seen in visual and motor systems (Benwell et al., 2014; Ikkai et al., 2016; Muret & Makin, 2021). These findings mirror homeostatic mechanisms—e.g., enhanced processing in an ignored visual field after unilateral deprivation (Lunghi et al., 2015; Turrigiano, 2012). The neurochemical basis of tRNS, particularly its effects on GABA and glutamate balance, may help explain the observed hemispheric differences, like previously studied with noninvasive brain stimulation (Stagg et al., 2009; Matsuta et al., 2022).

An interesting question arises from our findings: while the present study shows that unilateral tRNS over (right) frontoparietal areas can have inhibitory effects, previous research suggests that bilateral tRNS over the left and right IPS can enhance visual attention learning (Contò et al., 2021). Physiological studies indicate that electrical stimulation can produce opposite effects on neural activity, even with the same stimulation protocol, depending on the cortical state at the time of stimulation (Krause et al., 2022). One hypothesis is that bilateral tRNS enhances the overall functioning of the brain’s attentional network by maintaining a homeostatic balance between homologous areas in both hemispheres, thereby optimizing performance. In contrast, unilateral tRNS may disrupt this balance and prevent facilitation. However, this hypothesis remains to be empirically tested.

In conclusion, this study contributes to our understanding of hemispheric asymmetries in attentional control, showing how left and right frontoparietal tRNS differentially affect spatial attention across the visual field. These findings emphasize the importance of stimulation site, timing, and cortical state in shaping behavioral outcomes and lay the groundwork for future research and clinical translation.

